# Mutational signatures in upper tract urothelial carcinoma define etiologically distinct subtypes with prognostic relevance

**DOI:** 10.1101/735324

**Authors:** Xuesong Li, Huan Lu, Bao Guan, Zhengzheng Xu, Yue Shi, Juan Li, Wenwen Kong, Jin Liu, Dong Fang, Libo Liu, Qi Shen, Yanqing Gong, Shiming He, Qun He, Liqun Zhou, Weimin Ci

## Abstract

Parts of East Asia have a very high upper tract urothelial carcinoma (UTUC) prevalence, and an etiological link between the medicinal use of herbs containing aristolochic acid (AA) and UTUC has been established. The mutational signature of AA, which is characterized by a particular pattern of A:T to T:A transversions, can be detected by genome sequencing. Thus, integrating mutational signatures analysis with clinicopathological data may be a crucial step toward personalized treatment strategies for the disease. Therefore, we performed whole-genome sequencing (WGS) of 90 UTUC patients from China. Mutational signature analysis via nonnegative matrix factorization method defined three etiologically distinct subtypes with prognostic relevance: (i) AA, the typical AA mutational signature characterized by signature 22, had the highest tumor mutation burden, the best clinical outcomes; (ii) Age, an age-related group featured by signatures 1 and 5, had the lowest weighted genome instability index score, the worst clinical outcomes; and (iii) DSB, signature with deficiencies in DNA double strand break-repair featured by signatures 3, the intermediate clinical outcomes. Additionally, the distinct AA subtype was associated with AA exposure, the highest number of predicted neoantigens and heavier lymphocytes infiltrating. Thus, it may be good candidate for immune checkpoint blockade therapy. Notably, we showed AA-mutational signature was also identified in histologically “normal” urothelial cells. Thus, non-invasive urine test based on the AA-mutational signature could take advantage of this “field effect” to screen individuals at increased risk of recurrence due to exposure to herbal remedies containing the carcinogen AA. Collectively, the findings here may accelerate the development of novel prognostic markers and personalized therapeutic approaches for UTUC patients in China.

## Introduction

The epidemiology and clinicopathological characteristics of UTUC is largely influenced by its geographic distribution. Parts of East Asia such as Taiwan have a much higher UTUC prevalence, accounting for more than 30% of urothelial cancers (UCs)(1–3) compared to only 5-10% of UCs in Western populations(4). Chinese patients are more frequently female, of higher rate for kidney failure(5). Conceivably, geographic differences in risk factors for UTUC, such as AA-containing herb drugs consumption may account for some of these observations(1, 6–8). It has been shown that AA-induced mutational signature was characterized by A:T to T:A transversions at the sequence motif A[C | T]AGG(8, 9). The unusual genome-wide AA signature, termed signature 22 in COSMIC, holds great potential as “molecular fingerprints” for AA exposure in multiple cancer types(7, 9). Similarly, other endogenous or exogenous DNA damaging agents will also create mutational signatures detectable by DNA sequencing(10). Thus, analyses of mutational signatures, informative in other tumors(11, 12), have not been comprehensively reported in UTUC especially Chinese patients. The most comprehensive genomic studies of UTUC in Western populations used exome sequencing or targeted panel sequencing of cancer-associated genes(13–15). However, deriving signatures from whole-exome data may be problematic(16). Herein, we reported the first comprehensive genomic analysis of UTUC based on mutational signatures using WGS in 90 Chinese UTUC patients. Mutational signature analysis defined three UTUC subtypes with distinct etiologies and clinical outcomes. We further showed that geographically distinct AA subtype with high tumor mutation burden and higher level of tumor-infiltrating lymphocytes held great potential for immunotherapy. These results provided deeper insights into the biology of this cancer.

## Results

### Copy number variations (CNVs) dominate the UTUC landscape

The flowchart of selecting patients was shown in fig. S1. Clinicopathologic characteristics of 90 UTUC patients were listed in table S1. The samples were sequenced by WGS to an average 30X depth which allowed us to comprehensively catalog both single nucleotide variations (SNVs), small insertions and deletions (indels) and copy number variations (CNVs). We identified a median of 19,639 (interquartile range [IQR], 16,578 to 32,937) SNVs and a median of 2,197 (IQR, 1,615 to 2,650) indels. A median of 437 (IQR, 355 to 584) coding mutations was noted (fig. 1A). The increased mutation load was consistent with the increased prevalence of exposure to the potent mutagen AAs (fig. 1B). Next, we examined the candidate driver genes with MutSigCV (Q<0.05). Only two genes, TP53 and FRG2C, were identified (fig. 1C). However, many genes listed in the Cancer Gene Census as known driver genes were affected by non-silent mutations including genes which frequently mutated in Western UTUC patients, such as KMT2D (18%) and ARID1A (14%) (fig. 1C). Additionally, the overall genomic landscape confirmed the dominance of recurrent CNVs compared with SNVs and indels in the cohort (fig. 1D).

**Fig. 1.**
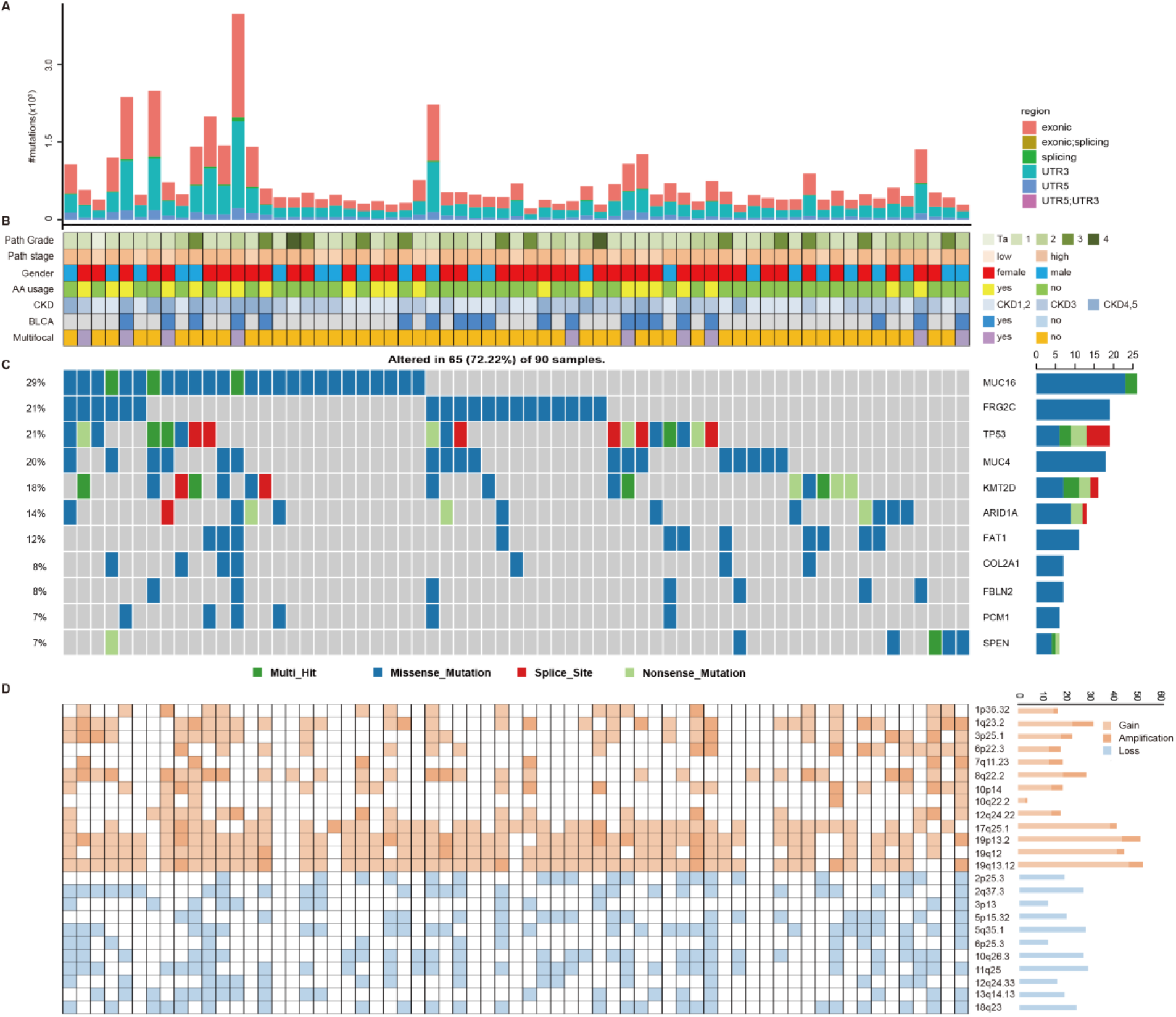
Oncoplot of genomic alterations in UTUC. **(A)** The number of coding mutations were represented by barplots. **(B)** Selected patient clinical features were included. **(C)** Cancer consensus frequently mutated genes and mutation types were indicated in the legend. **(D)** Selected loci with high recurrence rate among patients were shown. Copy number gain/loss and amplification/deletion were inferred as genes with GISTIC scores of 1 and 2, respectively. BLCA: synchronous bladder cancer, bladder recurrence and bladder cancer history. AA: aristolochic acid. CKD: chronic kidney disease.

### Mutational signatures define three UTUC subtypes with distinct etiology

Next, we explored the dynamic interplay of risk factors and cellular processes using mutational signature analysis. We identified 23 mutational signatures defined by COSMIC in our cohort, which were merged by shared aetiologies into 9 signatures by MutationalPatterns(17) (Materials and Methods). Hierarchical clustering based on the number of SNVs attributable to each signature confirmed three major subtypes (fig. 2A): (1) AA, there is significantly higher number of signature 22(1, 8) mutations (a median of 1,871) compared to the other two subtypes (fig. 2B, Wilcoxon rank test); (2) Age, there is significantly higher number of signatures 1 and 5, attributed to clocklike mutational processes accumulated over cell divisions(18) (fig. 2C, Wilcoxon rank test); and (3) DSB, there is significantly higher number of signature 3 mutations, associated with failure of DNA double-strand break-repair by homologous recombination (fig. 2D, Wilcoxon rank test). We further verified whether the subtypes defined by mutational signatures associated with their attributed etiological and clinicopathologic features. Consistent with previous epidemiological studies in Asian patients(6, 19–21), we found that the AA subtype was significantly associated with AA-containing herb drugs intake, poor renal function, female sex, multifocality and lower T stage (fig. 2E, Kruskal-Wallis test). Thus, the AA subtype defined by mutational signatures did capture the etiological subgroup of UTUC patients who had AAs exposure. Furthermore, consistent with the scenario that DNA damage repair deficiency may increase genomic instability, the DSB subtype exhibited both higher level weighted genome integrity index (wGII) and higher level microsatellite instability (MSI) compared to the Age subtype (fig. 2F and 2G). Additionally, lymphovascular invasion, an independent predictor of clinical outcomes for UTUC(22), was higher in the Age subtype compared to the DSB subtype. Although statistically significance was not achieved which may be due to limited number of cases (table S2).

**Fig. 2.**
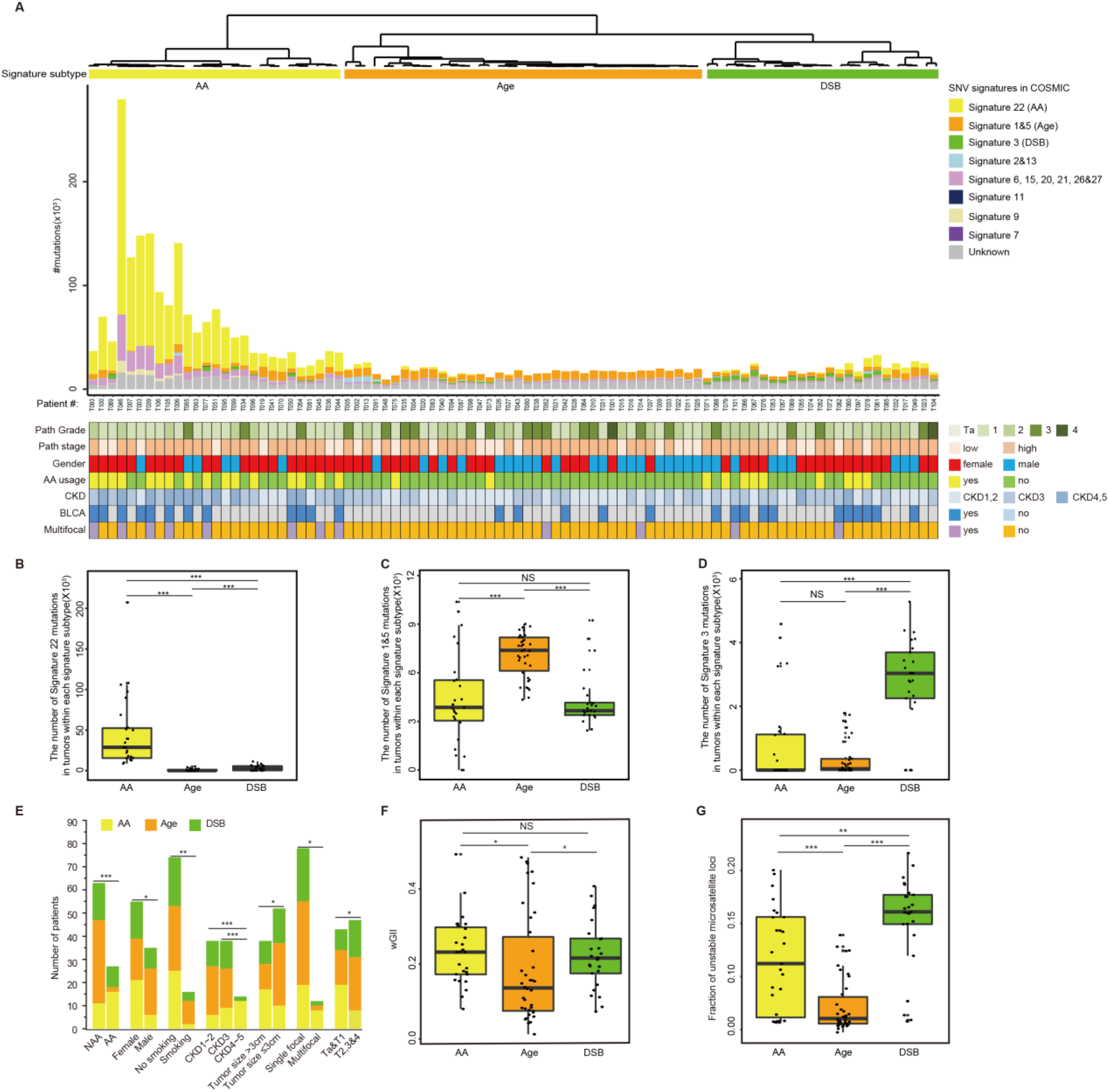
Genomic subtypes defined by mutational signatures. **(A)** Bar plot of the number of SNVs attributable to 9 merged signatures in each of the 90 tumors, sorted by hierarchical clustering (dendrogram at bottom), revealing AA-related (AA, yellow), Age-related (Age, orange), and DNA damage repair deficient (DSB, green). Selected clinical features were represented in the bottom tracks. **(B)** The box plot showed the number of signature 22 mutations in tumors within each signature subtype. \**p*<0.05; \*\**p*<0.01; \*\*\**p*< 0.001 (Wilcoxon rank test). **(C)** The box plot showed the number of signatures 1&5 mutations in tumors within each signature subtype. \**p*<0.05; \*\**p*<0.01; \*\*\**p*< 0.001 (Wilcoxon rank test). **(D)** The box plot showed the number of signature 3 mutations in tumors within each signature subtype. \**p*<0.05; \*\**p*<0.01; \*\*\**p*< 0.001 (Wilcoxon rank test). **(E)** The bar graph showed the association between three subtypes and clinicopathologic features. \**p*<0.05; \*\**p*<0.01; \*\*\**p*< 0.001 (Kruskal-Wallis test). **(F)** Comparison of the weighted Genome Instability Index (wGII) scores among three subtypes. **(G)** Microsatellite instability (MSI) status was evaluated by fraction of unstable microsatellite loci identified by WGS data. *P*-values were calculated by Wilcoxon rank test (\**p*<0.05, \*\**p*< 0.01, \*\*\**p*< 0.001).

### Subtypes defined by mutational signatures can predict clinical outcomes

Next, we explored whether clinical outcomes of the three subtypes differed significantly. Notably, a Kaplan-Meier plot revealed that the subtypes can predict both cancer-specific survival (CSS) (fig. 3A, log-rank, *p*=0.006) and metastasis-free survival (MFS) (fig. 3B, log-rank, *p*=0.007). The AA and DSB subtypes exhibited favourable outcomes compared with the Age subtype. Since an improved prognostic stratification could offer more tailored therapeutic decisions, we further stratified the patients into subgroups, muscle invasion and non-muscle invasion subgroups. Consistently, the AA and DSB subtypes also exhibited favourable outcomes in muscle-invasive UTUC patients (log-rank, *p*=0.001 for CSS, log-rank, *p*=0.002 for MFS) (fig. 3C and 3D). In multivariate Cox regression proportional analyses adjusted for the effects of standard clinicopathologic variables, mutational subtypes were still an independent risk factor for CSS (Hazard Ratio [HR]=4.05, 95% CI: 1.58~10.42, p=0.004) and MFS (HR=3.43, 95% CI: 1.46~8.06, p=0.006) (table S3). We also used the recently described mutational signature deconvolution method, Mutalisk, for comparison(23). This method was based on different statistical frameworks, and therefore some differences were to be expected. Nevertheless, we observed the same key signature patterns with similar-sized patient subgroups expressing the dominant signature types (fig. S2A). Both Mutalisk and MutationalPatterns identified the same group of patients with AA signature. MutationalPatterns identified 9 putative Age subtype of UTUC patients that Mutalisk did not identify (T002, T005, T013, T040, T047, T048, T087, T091 and T094) (fig. S2B). Since the Age subtypes by either MutationalPatterns and Mutalisk showed the worst CSS and MFS (fig. 3, fig. S2C and S2D), we would propose that MutationalPatterns was preferable to err on the side of overcalling rather than undercalling the presence of age-related signature.

**Fig. 3.**
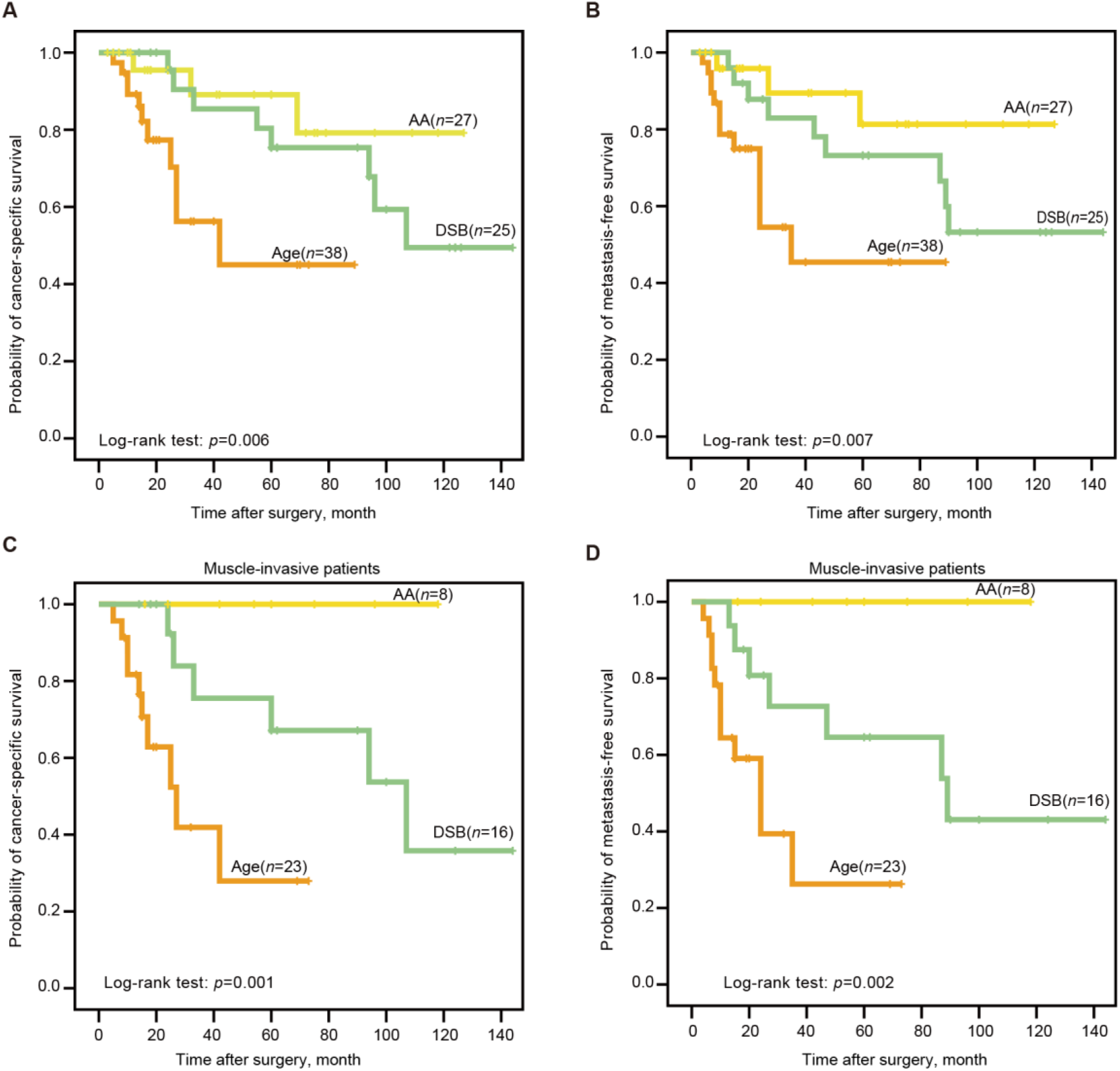
Subtypes defined by mutational signatures predict clinical outcome. **(A)-(D)**Kaplan-Meier survival curves showed the mutational signature subtypes can predict both CSS and MFS for the whole cohort, as well as for muscle-invasive UTUC patients. CSS: cancer-specific survival. MFS: metastasis-free survival. *P*-values were calculated by the log-rank test. *n*, the number of cases.

### Integration of mutational signature subtypes with copy number alterations

One practical question was how this subtyping could be implemented clinically. Previous study had showed that 10X coverage data can confidently retrieve dominant signatures(11). Therefore, lower-coverage WGS could provide a cost-effective alternative for signature-based stratification especially for the AA subtype (more than 35% mutations contributed by signature 22 in our cohort). Although for the Age and DSB subtype which did not have the dominant signatures, we next explored whether CNV features can distinguish these two subtypes. First, we examined the landscape of CNVs in UTUC by ichorCNA(24) which can predict CNVs in tumor-only samples. Recurrent CNVs in UTUC patients of BCM/MDACC cohort were also identified in our study(13), such as gain in 1q, 8q and loss in 2q, 5q, 10q (fig. S3A). However, the defined mutational subtypes presented similarities in terms of CNV profiles (fig. S4A and S4B). Of note, several focal CNVs were significantly associated with the mutational subtypes, such as deletion of 10q26.2-26.3 was more prevalent in the DSB subtype compared to the Age subtype, and amplification of 1q31.1-1q32.1 was more prevalent in the AA subtype compared to the DSB subtype (fig. S4C). Moreover, we performed unsupervised hierarchical clustering which separated UTUCs into three groups with distinct CNV patterns. However, there was not significantly association between mutational signature subtypes and CNV subtypes (fig. S5A). Meanwhile, the overall survival of the CNV subtypes did not differ significantly (fig. S5B and S5C). Among all the recurrent CNV regions, only amplification of 17q25.1 was significantly associated with better overall survival (fig. S5D). Hence, the CNV profiles seem to capture a different type of clinicopathological information compared with the mutational signature subtypes.

### Field cancerization may contribute to malignant transformation especially for the AA subtype

Clinical interest in the AA subtype is increasing especially in East Asia since AA-associated UTUCs are prevalent(1, 25). Consistent with our previous epidemiological study(21), we found an increased rate of multifocality and high bladder recurrence in the AA subtype (fig. 2B). Field cancerization(26), which is the development of a field with genetically altered cells, has been proposed to explain the development of multiple primary tumours and local recurrence. Therefore, we further sequenced three AA subtype patients (table S4) including a multifocal patient (fig. 4A and fig. S6). We did find that significant amount of SNVs and indels in the morphologically normal urothelium specimens in all three patients. Strikingly, the AA mutational signature was consistently identified in urothelium specimens and tumor tissues in all 3 patients (fig. 4B), which indicated that AA exposure may contribute to the field cancerization. Probably damaging mutations in genes that were documented by the COSMIC and the KEGG pathway in cancer were shown in table S5. Moreover, in the multifocal patient, the urothelial tumour at renal pelvic from 2007 shared no genetic alterations with a renal pelvic tumour from 2015 or a bladder tumour from 2015. However, the two tumours from 2015 were genetically related (fig. 4C). Therefore, field cancerization and intraluminal seeding could co-contribute to the multifocality and increased bladder recurrence in the AA subtype patients.

**Fig. 4.**
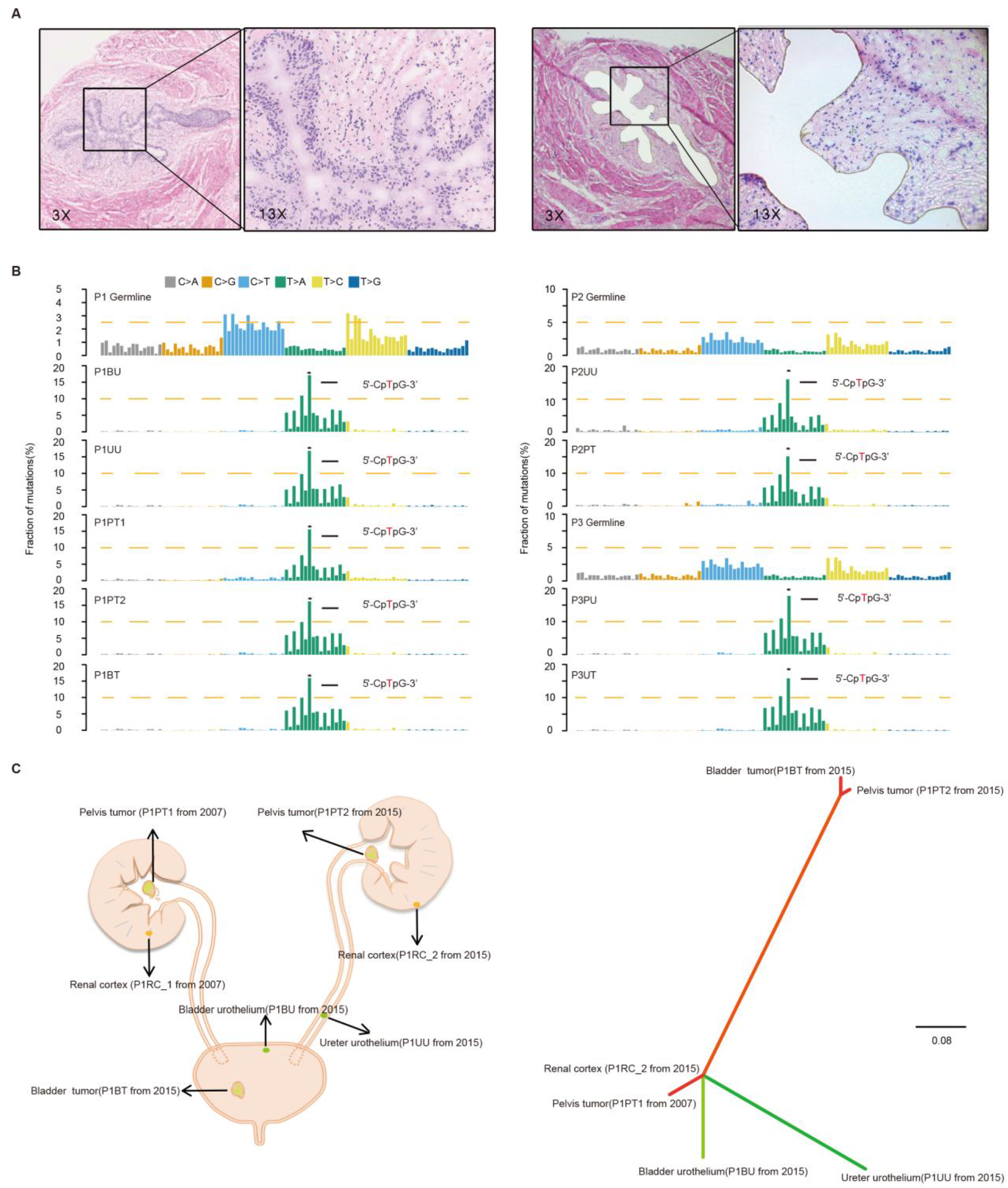
Field cancerization may contribute to malignant transformation especially for the AA subtype. **(A)** Laser capture microdissection of urothelium from a representative patient. Magnified insets (13x) of the black frame area showed ureteral urothelium before laser capture and the remaining cells after laser capture. **(B)** Trinucleotide contexts for somatic mutations in biopsies from three patients of the AA subtype. The height of each bar (the *y*-axis) represented the proportion of all observed mutations that fall into a particular trinucleotide mutational class. AA: aristolochic acid; UU: ureter urothelium; BU: bladder urothelium; RC: renal cortex; BT: bladder tumor; UT: urothelium tumor; and PT: pelvis tumour. **(C)** Phylogenetic relationships of the six samples from the multifocal patient were deciphered using mrbayes_3.2.2. Spatial locations of core biopsies of the patient were presented in the left panel.

### Potential for immunotherapy in the AA subtype of UTUC patients

The AA subtype bears high mutation burdens thus may be good candidate for immune checkpoint blockade therapy(27). To address this possibility, we predicted neoantigens binding to patient-specific human leukocyte antigen (HLA) types. The AA subtype had the highest number of predicted neoantigens (fig. 5A). Moreover, it has been reported that lymphocytes infiltration especially CD3+ lymphocytes in tumour region is associated with improved survival in a range of cancers including urothelial cancer(28–30). Previous study has shown that the number of tumor-infiltrating lymphocyte independently correlates progress-free survival in NSCLC patients treated with nivolumab immunotherapy(31). Pathological assessment of tumor-infiltrating lymphocytes allows an easy evaluation of immune infiltration. Therefore, we further evaluated the extent of tumor-infiltrating mononuclear cells (TIMCs) and CD3+ lymphocytes in 76 available samples (table S6). We found the number of CD3+ lymphocytes was positively associated with the number of stromal TIMCs (R^2^=0.72; *p*<0.001) (fig. 5B). The AA subtype had the highest number of both stromal TIMCs (Wilcoxon rank test, *p*<0.001) and CD3+ lymphocytes (Wilcoxon rank, *p*<0.001) (fig. 5C and 5D). Representative images from a patient of the AA subtype and the Age subtype were shown in the fig. 5E and fig. 5F, respectively.

**Fig. 5.**
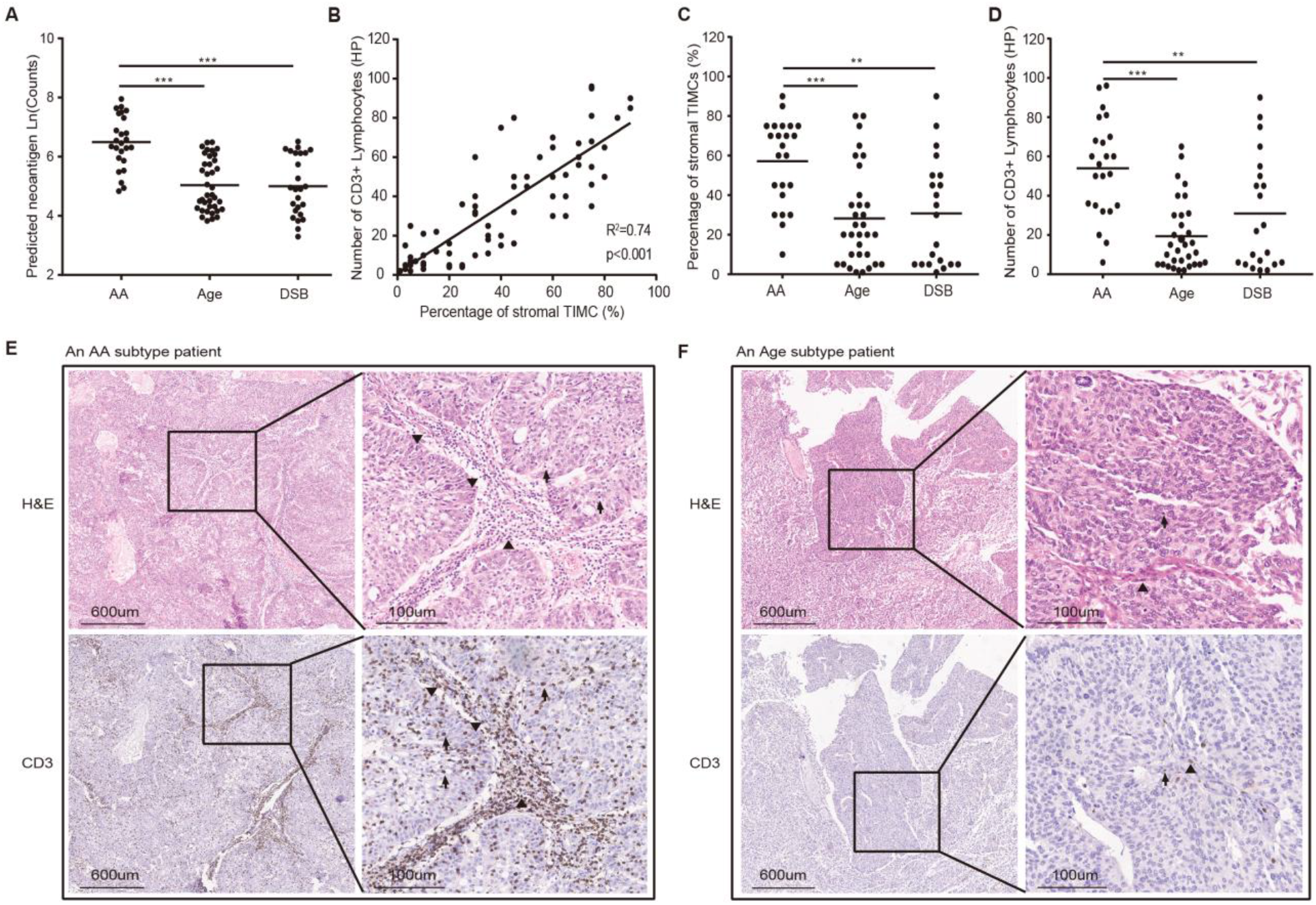
Predicted neoantigens and tumor-infiltrating lymphocytes in each mutational subtype. **(A)** Neoantigen burden was significantly higher in the AA subtype. Statistical significance was determined by Wilcoxon rank test (\*\*\**p*<0.001). **(B)** Positive correlation of the percentage of stromal tumor-infiltrating mononuclear cells (TIMCs) and the number of CD3+ lymphocytes in 76 UTUC patients of our cohort (nAA=23; nAge=32; nDSB=21). The percentage of stromal TIMCs **(C)** and CD3+ lymphocytes **(D)** were shown in each subtype of patients. Statistical significance was determined by the Wilcoxon rank test (\*\**p*<0.01, \*\*\**p*<0.001). HP represented high-power field. Images of TIMCs and CD3+ lymphocytes of a representative patient from the AA subtype **(E)** and the Age subtype **(F)** Triangle highlighted the TIMCs or CD3+ lymphocytes in stromal tumour region. Arrow highlighted the TIMCs or CD3+ lymphocytes in intra-tumour region.

## Discussion

Herein, we performed the first comprehensive genomic analysis of Chinese UTUC patients. Three subtypes were defined by mutational signatures. We found that the AA subtype was significantly associated with the usage of AA-containing herb drugs. However, a substantial percentage of patients with the AA-mutational signature did not report usage of AA-containing herb drugs. This finding may be due to the difficulty to track the dosage of AAs from various herbal remedies. Some AA-containing herbal remedies have been officially prohibited in China since 2003, but Aristolochia contains over 500 species, many of which have medicinal properties(32). In the current study, we only took 70 AA-containing single products, such as *Guang Fangchi, Qing Mu Xiang, Ma Dou Ling, Tian Xian Teng, Xun Gu Feng*, and *Zhu Sha Lian* in Mandarin Chinese and mixed herbal formulas, such as, *Long Dan Xie Gan*, into account. And the detail list of AA-containing herb drugs surveyed by current study was shown in table S7. Some herb plants containing AA were not banned such as *Xi Xin(33)*. Exposure to AAs and their derivatives may still be widespread in China. Notably, we showed AA-mutational signature was also identified in histologically “normal” urothelial cells. In a published study(34), similar “field effect” was also identified in an Chinese UTUC patient. Thus, non-invasive urine test based on the AA-mutational signature could take advantage of this “field effect” to screen individuals at increased risk of recurrence due to exposing to herbal remedies containing the carcinogen AA.

One limitation of current study was lack of a matched normal for 85 of 90 UTUC patients due to lack of funds. Therefore, we relied on the public mutation database and in-house genomic data of 1000 healthy Chinese individuals for the filtering germline variances. But in recent years, these catalogs of human variation have grown exponentially, questioning the necessity of sequencing a matched normal for every tumor sample(35). Additionally, a matched normal will not be available in many pathological conditions, such as cancer cell lines and liquid biopsy. Although we can not make definite conclusions that the SNVs identified in current study were somatic mutations. We may also overcall the Age subtypes without a matched normal since germline variants contributed to signature 5 in COSMIC which combined with signature 1 as age-related signature in current study. However, with this caveat, the mutational signature subtypes did capture distinct clinical and genomic features, such as better overall survival, higher level of wGII and higher level of MSI in DSB subtype compared to Age subtype of patients.

Additionally, our study supports a rationale for management of UTUC, most notably the AA subtype. Patients with AA mutational signature may predict better checkpoint inhibitors response. Strikingly, the DSB subtype was featured by signature 3 in COSMIC. In pancreatic cancer, responders to platinum therapy usually exhibit signature 3 mutations(36). These results suggested that platinum-based therapy may also benefit UTUC patients of DSB subtype. More recently, protein components of the DNA damage repair (DDR) systems have been identified as promising avenues for targeted cancer therapeutics(37). We also further assessed the mutation rates across 345 genes associated with DDR, as previously described in a pan-cancer analysis(37) (fig. S7). There was a 1.6 fold enrichment (P=0.007, 95% CI:1.12-2.20) and a 1.4 fold enrichment (P=0.01, 95% CI:1.07-1.86) of DDR genes alterations in DSB and Age subtype, respectively. Further mechanistic insights of the DSB and Age subtypes may provide the basis for therapeutic opportunities within the DNA damage response. In summary, our study provides the most comprehensive genomic profile of Chinese UTUC patients to date. The findings may accelerate the development of novel prognostic markers and personalized therapeutic approaches for UTUC patients especially those from geographical regions with exposure to AA-containing herb drugs.

## Materials and Methods

### Patients cohort

All the fresh samples in this study were obtained from Peking University First Hospital between January 2016 to December 2017 (Grant No.2015[977]). These fresh samples were stored in liquid nitrogen after surgery immediately. Formalin-fixed, paraffin-embedded (FFPE) samples were provided by the Institute of Urology after pathologic diagnosis. The main endpoint events consisted of overall survival (OS), cancer-specific survival (CSS), metastasis-free survival (MFS), progression-free survival (PFS) and bladder-recurrence free survival (BRFS). All patients were not treated with neoadjuvant chemotherapy. AA exposure assessment was performed according to self-reported data on 70 AA and its derivatives-containing herb drugs (collectively, AA) intake(21, 38). These herbs were taken as single products (*Guan Mu Tong (Aristolochia manshuriensis Kom), Guang Fangchi (Aristolochia fangchi), Qing Mu Xiang (Radix Aristolochiae), Ma Dou Ling (Fructus Aristolochiae), Tian Xian Teng (Caulis Aristolochiae), Xun Gu Feng (herba Aristolochiae mollissimae*), and *Zhu Sha Lian* (*Aristolochia cinnabarina*)) or were components of mixed herbal formulas (eg, *Guan Mu Tong* in the *Long Dan Xie Gan* mixture). And the detail list of AA-containing herb drugs surveyed by current study was shown in table S7. The accumulated self-reported usage for the above drugs more than a year was termed as AA exposure patients. Clinical and demographic information was obtained from a prospectively maintained institutional database. The study was approved by the Ethics Committee of Peking University First Hospital.

### Whole-genome sequencing

The sheared DNA was repaired and 3’ dA-tailed using the NEBNext Ultra II End Repair/dA-Tailing Module unit and then ligated to paired-end (PE) adaptors using the NEBNext Ultra II Ligation Module unit. After purification by AMPure XP beads, the DNA fragments were amplified by PCR for 6-8 cycles. Finally, the libraries were sequenced with an Illumina X-ten instrument, thus generating 2 x 150-bp PE reads.

### Single nucleotide variations (SNVs) and indels calling

SNVs and indels were called using VarScan2(39)and Vardict(40) in the form of single sample mode, then Rtg-tools(41) was used to remove variants called in a set of 1000 healthy, unrelated Chinese individuals (unpublished data from CAS Precision Medicine Initiative) and got the common variants called by the two software. Further filtering criteria were carried out according to the reference(11, 42). Finally, Annovar(43) was applied to remove variants whose mutation frequency was no less than 0.001 in 1000 Genomes project, latest Exome Aggregation Consortium dataset, NHLBI-ESP project with 6500 exomes, latest Haplotype Reference Consortium database and latest Kaviar database.

### Mutational signature analysis

R package MutationalPatterns(17) was carried out to assign each sample’ signature. We discovered 23 COSMIC signatures which were merged by shared etiologies into 9 signatures in our cohort. We named signature 22 which has been found in cancer samples with known exposures to aristolochic acid(1, 8) as AA, DNA double strand break-repair deficiency featured by signature 3 as DSB. We merged signature 1 and 5 as Age(18), signature 2, 13 as APOBEC, combined signatures including signature 6, 15, 20, 21, 26, 27 as MSI, combined signatures including signature 8, 12, 16, 17, 18, 19, 23, 25, 28, 30 as unknown. Mutational signatures was extracted from the mutation count matrix, with non-negative matrix factorization (NMF). Hierarchical clustering was performed by the number of single nucleotide variants (SNVs) attributable to each signature.

### Copy number variations (CNVs) analysis

To determine copy number states, we used a recently published method(44). Briefly, the genome was divided into bins of variable length with an equal amount of potential uniquely mapping reads. We simulated approximately 37X hg19 reads mapped by BWA and divided the genome into 50,000 bins of the same mapping ability. Loess normalization was used to correct for GC bias.

### Candidate driver gene analysis

Candidate driver genes were explored by MutSigCV(45) to analyse SNVs and indels. Chromatin state, DNA replication time, expression level and the first 20 principal components of 169 chromatin marks(46) from the RoadMap Epigenomics Project(47) of the mutated gene were used as covariates. We applied the cut off 0.05 for Q value. Cancer census genes from COSMIC(48) release v87 were calculated for their mutation rates (mutations/gene length) and genes with more than 10 samples were selected with high mutation rates.

### The weighted genome integrity index score (wGII) analysis

We calculated the wGII score as reported(49). Briefly, the percentage of gained and lost genomic material was calculated relative to the ploidy of the sample. The use of percentages eliminates the bias induced by differing chromosome sizes. The wGII score of a sample was defined as the average of this percentage value over the 22 autosomal chromosomes.

### Microsatellite instability (MSI) analysis

mSINGS(50) was used to calculate the ratio of unstable loci in the genome of each sample. First, the microsatellites at the reference genome were calculated by MSIsensor(51). Sixteen microsatellite stable esophageal squamous cell carcinomas(52) were used to calculate the baseline for the absence of MSI signature (signature 6, 15, 20, 21, 26, 27) as defined by COSMIC(48).

### Neoantigen prediction

HLAscan(53) was used to genotype the HLA region with HLA-A, HLA-B and HLA-C taken into consideration. NetMHC4.0(54) was used for predictions of peptide-MHC class I interaction, SNVs annotated by Annovar(43) as nonsynonymous were used to do this analysis. An in-house script was carried out to obtain possible 9–amino acid sequences covering the mutated amino acids according to the manual. Rank in the output that was no more than two were considered as binding in the light of BindLevel suggestion.

### Assessing tumor-infiltrating lymphocytes

The tumor infiltrating mononuclear lymphocytes were measured according to a standardized method from the International Immuno-Oncology Biomarkers Working Group(55). And the CD3 antibody (ab5690, 1:10000; Abcam) was used to evaluate the CD3+ lymphocytes in tumor section.

## Supporting information

Main text_2019.pdf

## Acknowledgements

We thank Gengyan Xiong and Lei Zhang for scientific inputs about the manuscript as well as Han Hao, Qin Tang and Xiaoteng Yu for assistance with clinicopathological characterizations.

## Funding

This work was supported by CAS Strategic Priority Research Program (XDA16010102 to W.C.), the National Key R&D Program of China (2018YFC2000100 to W.C.), CAS (QYZDB-SSW-SMC039 to W.C.), the National Natural Science Foundation of China (7152146 to X.L., 81672541 to W.C. and 81672546 to L.Z.), K.C. Wong Education Foundation to W.C., the Clinical Features Research of Capital (Z151100004015173 to L.Z.), the Capital Health Research and Development of Special (2016-1-4077 to L.Z.).

## Disclosure

Weimin Ci certifies that there are no conflicts of interest.

## Data availability

The raw sequence data reported in this paper have been deposited in the Genome Sequence Archive(56) in BIG Data Center(57), Beijing Institute of Genomics (BIG), Chinese Academy of Sciences (accession numbers HRA000029). That can be accessed at http://bigd.big.ac.cn/gsa-human/s/HiObV4f3.

## Notes

https://bigd.big.ac.cn/gsa-human/s/HiObV4f3

