## Supplementary material for "Mutational signatures in upper tract urothelial carcinoma define etiologically distinct subtypes with prognostic relevance": Main text_2019.pdf

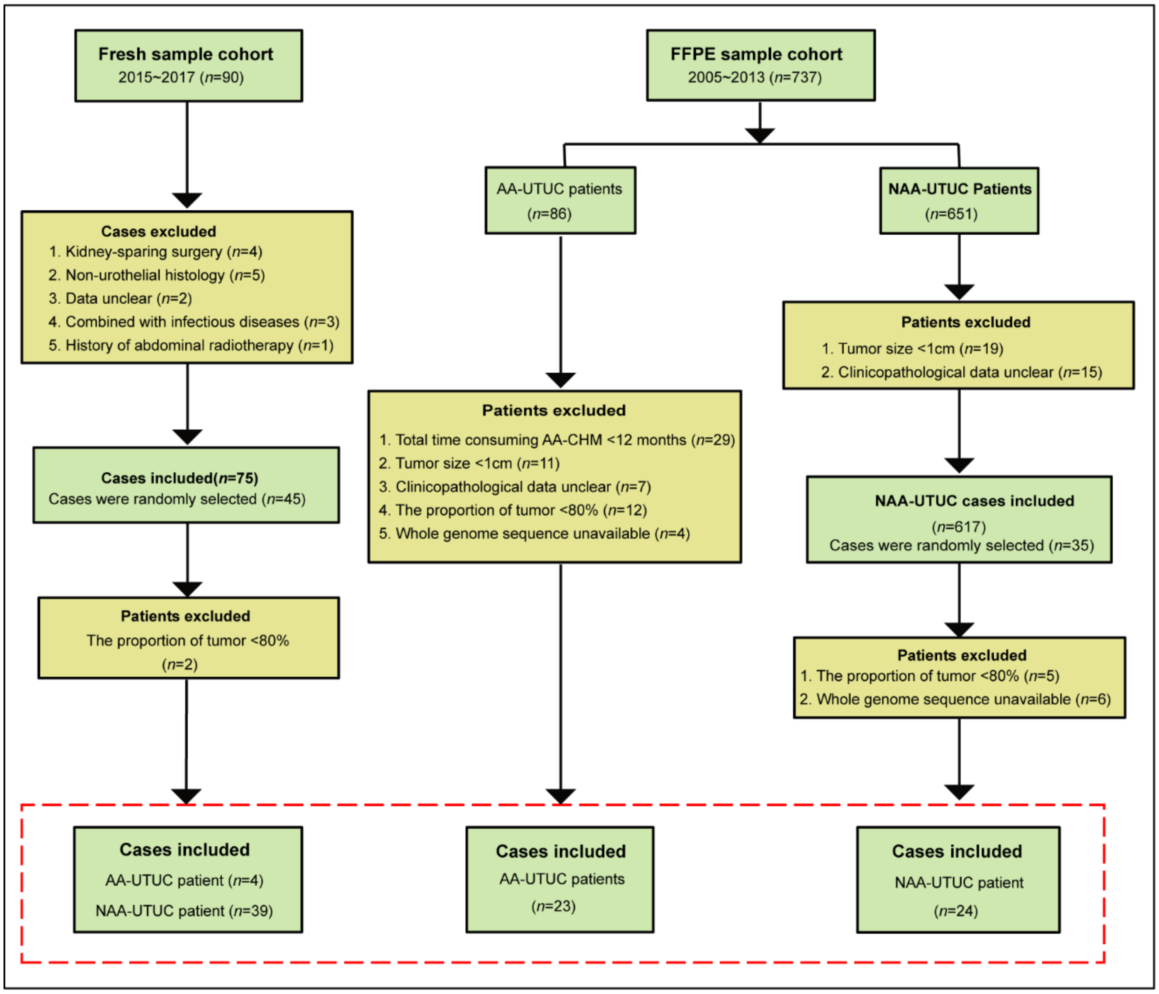


**Fig. S1. Flow chart of the patient selection.** The quality assurance review of clinicopathologic data was assessed by two senior pathologists. None of patients received neoadjuvant treatment. RNU: Radical nephroureterectomy, AA: aristolochic acid, FFPE: formalin-fixed paraffin-embedded.

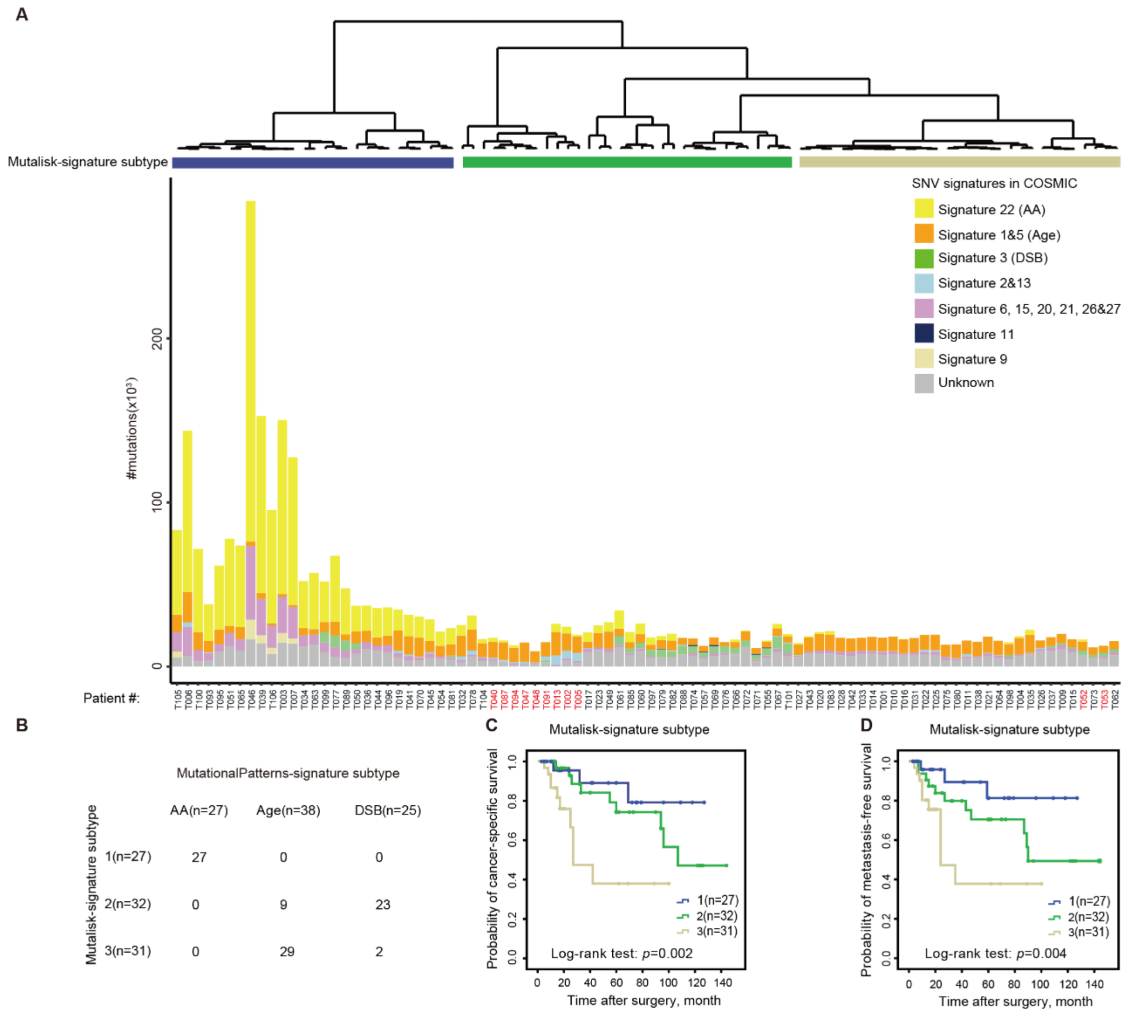


**Fig. S2**. **The mutational signature subtypes deconvoluted by Mutalisk. (A)** Bar plot of the number of SNVs attributable to 8 merged signatures in each of the 90 tumors, sorted by hierarchical clustering (dendrogram at bottom), revealing type 1 (*n*=27), type 2 (*n*=32), and type 3 (*n*=31). **(B)** Both Mutalisk and MutationalPatterns identified the same key signature patterns with similar-sized patient subgroups expressing the distinctive signature types. MutationalPatterns identified 9 putative Age subtype of UTUC patients that Mutlisk did not identify (T002, T005, T013, T040, T047, T048, T087, T091 and T094). **(C)** and **(D)** Kaplan-Meier survival curves showed that the Mutalisk subtypes can also predict both CSS and MFS. CSS: cancer-specific survival, MFS: metastasis-free survival. *P*-values were calculated by the log-rank test. *n*, the number of cases.

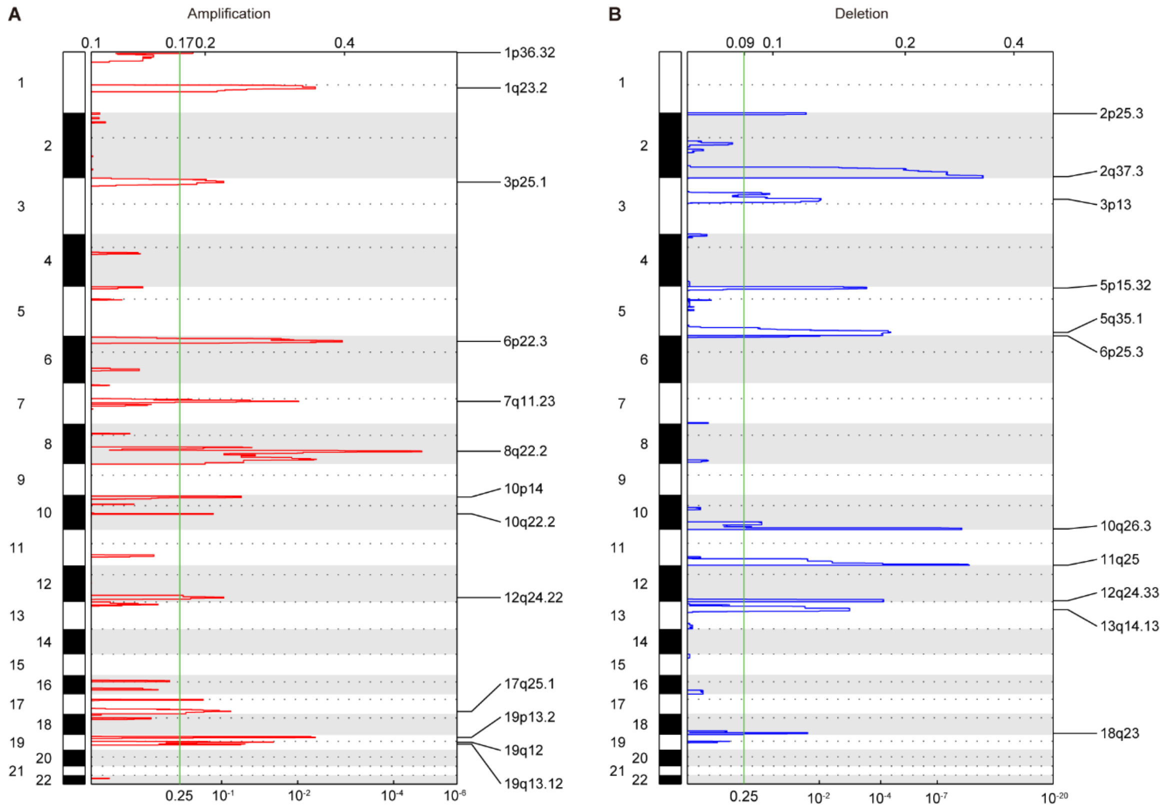


**Fig. S3. The CNV profiles in our cohort. (A)** and **(B)** GISTIC plot of the identified deletion and amplification peaks in the cohort.

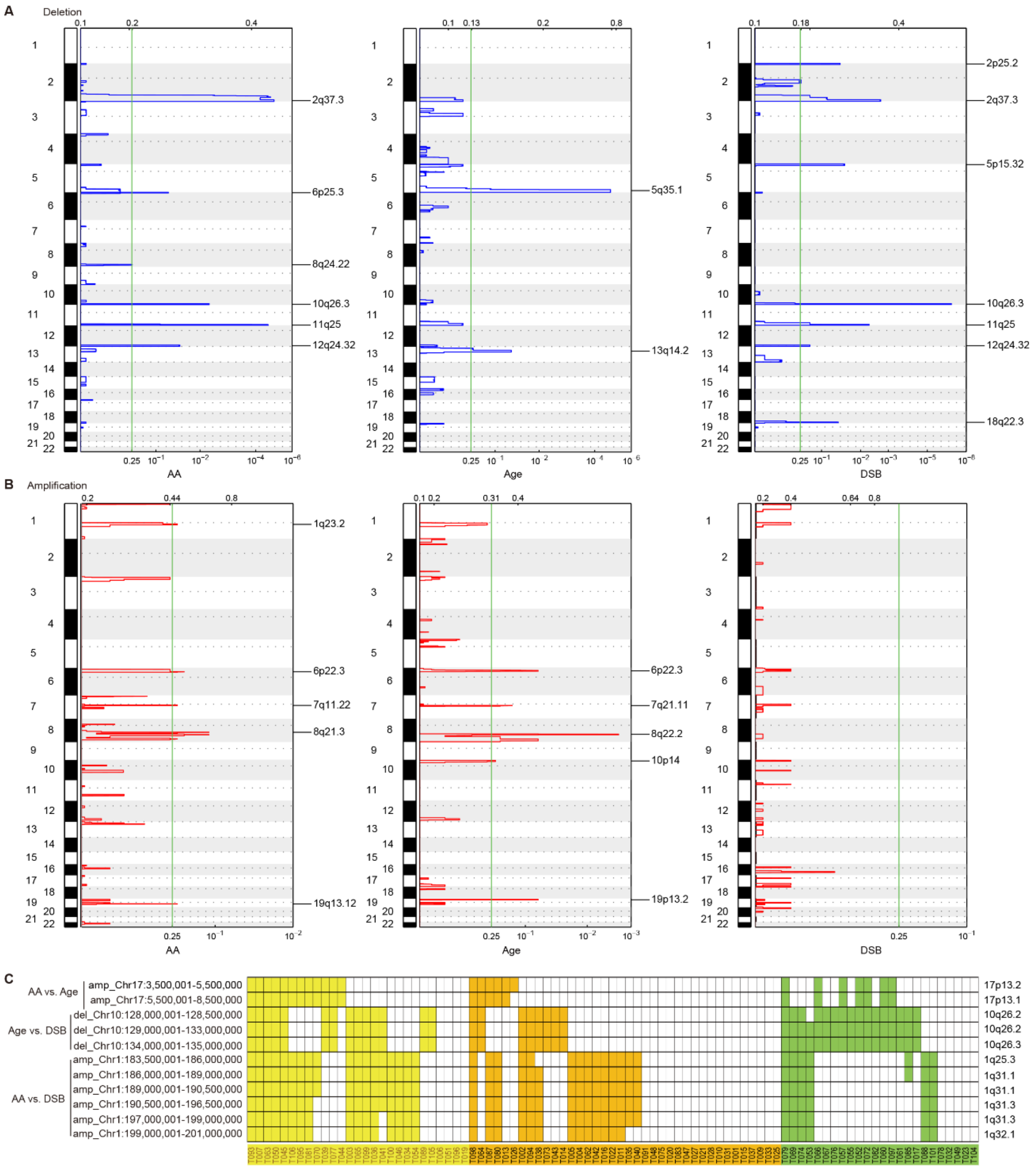


**Fig. S4. Integration of mutational signature subtypes with copy number alterations. (A)** and **(B)** GISTIC plots of the identified deletion and amplification peaks in the three mutational subtypes. **(C)** Several focal CNVs that were significantly associated with the mutational subtypes.

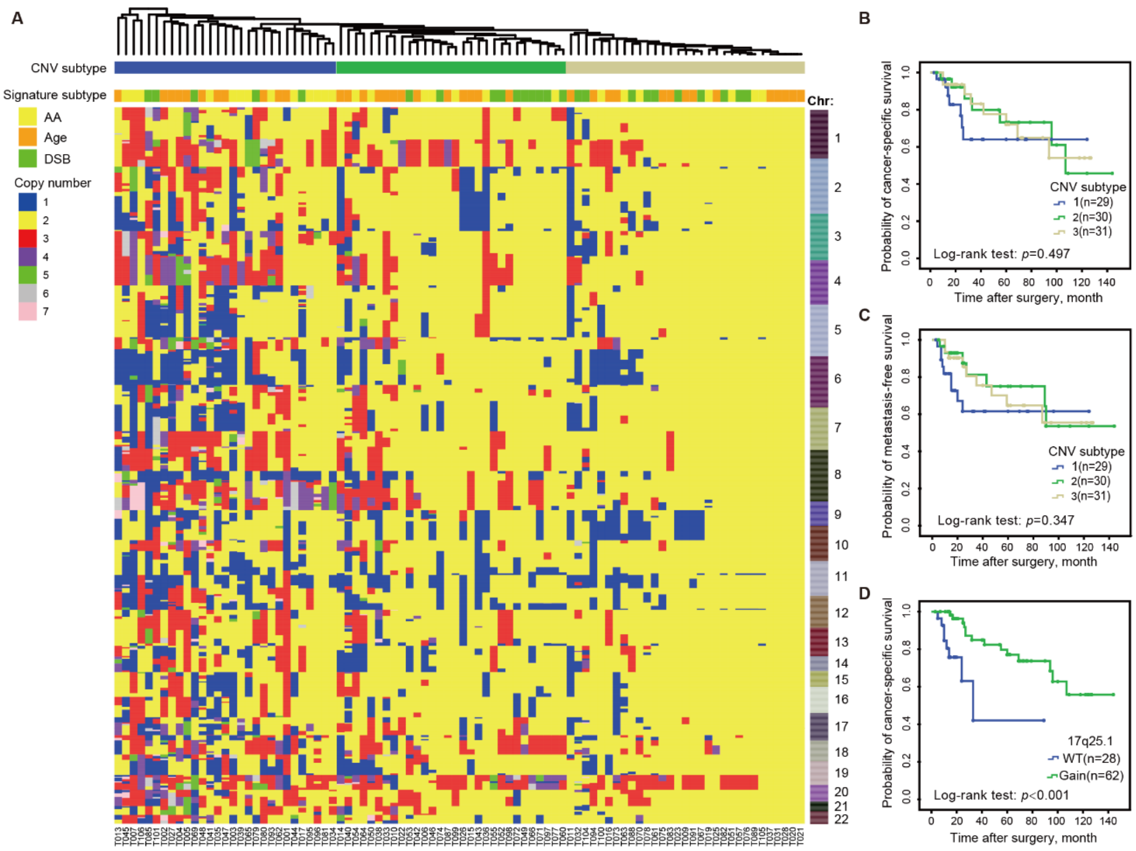


**Fig. S5. Genomic subtypes defined by CNV profiles. (A)** CNV profiles within 500k bin were evaluated by ichorCNA. The patients were sorted by hierarchical clustering (dendrogram attop), revealing type 1 ($n$=29), type 2 ($n$=30), and type 3 ($n$=31). The defined mutational signature type of the corresponding cases was indicated. **(B)** and **(C)** Kaplan-Meier survival curves showed that the CNV subtypes can not predict both CSS and MFS. CSS: cancer-specific survival. MFS: metastasis-free survival. $P$-values were calculated by the log-rank test. $n$, the number of cases. **(D)** Kaplan-Meier survival curves showed that only recurrent gain of 17q25.1 in our cohort can predict CSS. WT: wide type.

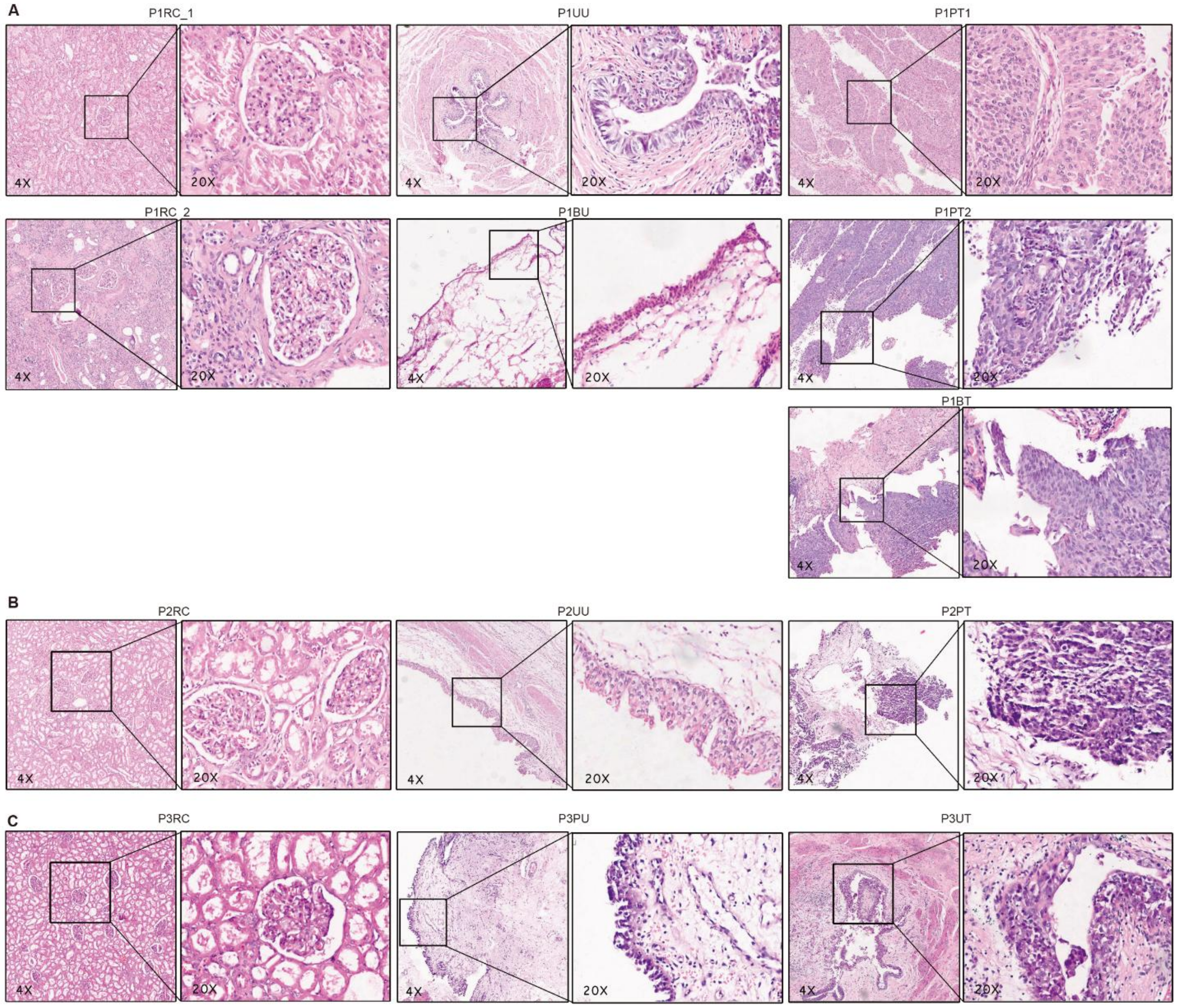


**Fig. S6. Histology of 13 sections from 3 UTUC patients within the AA subtype**. (A), (B) and (C), Urological histological sections in P1, P2 and P3 identified by H&E staining (4x). Insets represent magnification of the black frame area (20x). RC: renal cortex; PU: Pelvic urothelium; PT: renal pelvis tumour; UU: ureteral urothelium; UT: ureteral tumour; BU: bladder urothelium; BT: bladder tumour. Sample names were in the form of patient ID+sample type. For example, P3PU represented the pelvic urothelium from patient 3.

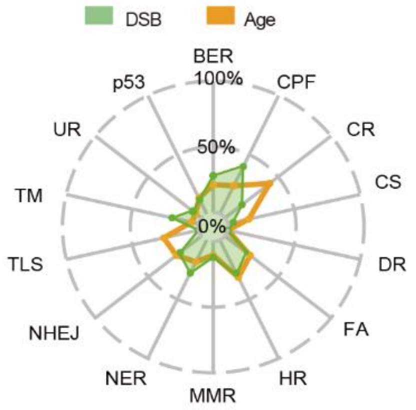


**Fig. S7. Mutated genes in DNA damage repair pathways of the DSB and Age subtypes, respectively.** The proportion of samples mutated in DNA damage repair pathways of the DSB and Age subtypes, respectively. AM, alternative mechanism for telomere maintenance; BER, base excision repair; CPF, checkpoint factor; CR, chromatin remodeling; CS, chromosome segregation; DR, direct repair; FA, Fanconi anaemia pathway; HR, homologous recombination; MMR, mismatch repair; NER, nucleotide excision repair; NHEJ, non-homologous end joining; TLS, translesion synthesis; TM, telomere maintenance; UR, ubiquitination response.

**Table S1. Clinicopathologic characteristics of 90 UTUC patients**

| Variable | Number (%) |
|---|---|
| Total | 90 |
| Median age at diagnosis, year (IQR) | 65 (60~71) |
| Gender | |
| Female | 55 (61.1%) |
| Male | 35 (38.9%) |
| Smoking | |
| Absent | 74 (82.2%) |
| Present | 16 (17.8%) |
| AA intake | |
| Absent | 63 (70.0%) |
| Present | 27 (30.0%) |
| Synchronous bladder cancer | |
| Absent | 85 (94.4%) |
| Present | 5 (5.6%) |
| History of bladder cancer | |
| Absent | 81 (90.0%) |
| Present | 9 (10.0%) |
| CKD | |
| 1 | 9 (10.0%) |
| 2 | 29 (32.2%) |
| 3 | 38 (42.2%) |
| 4 | 2 (2.2%) |
| 5 | 12 (13.2%) |
| Location | |
| Pelvis | 58 (64.4%) |
| Ureter | 32 (35.6%) |

| | |
|---|---|
| Multifocality | |
| Absent | 78 (86.7%) |
| Present | 12 (13.3%) |
| Size | |
| <3 cm | 38 (42.2%) |
| ≥3 cm | 52 (57.8%) |
| Architecture | |
| Papillary | 64 (71.1%) |
| Sessile | 26 (28.9%) |
| T stage | |
| Ta | 1 (1.1%) |
| T1 | 42 (46.7%) |
| T2 | 25 (27.8%) |
| T3 | 20 (22.2%) |
| T4 | 2 (2.2%) |
| G | |
| Low | 25 (27.8%) |
| High | 65 (72.2%) |
| N stage | |
| N0 or Nx | 83 (92.2%) |
| N1 | 5 (7.8%) |
| N2 | 2 (2.2%) |
| LVI | |
| Absent | 82 (8.9%) |
| Present | 8 (91.1%) |
| Postoperative chemotherapy | 9 (10.0%) |
| Postoperative radiotherapy | 5 (5.6%) |
| Survival data | |
| Cancer-related death | 22 (24.4%) |
| Overall survival | 24 (26.7%) |
| Metastasis | 25 (27.8%) |
| Bladder recurrence | 19 (21.1%) |
| Cancer progression | 39 (43.3%) |
| Median FU(IQR) | 25 (14~69) |
| Median FU for surviving patients, month(IQR) | 24 (14~70) |

Nx: No lymph node dissection was performed; IQR: interquartile range; LVI: lymphovascular invasion. FU=follow-up; NA=not applicable. AA=aristolochic acid; CKD=chronic kidney disease. All patients were not treated with neoadjuvant chemotherapy.

**Table S2. Clinical characteristics stratified by mutational signature**

| **Variable** | **No. (%)** | **AA (%)** | **Age (%)** | **DSB (%)** | ***P*-value** |
|---|---|---|---|---|---|
| Total | 90 | 27 | 38 | 25 | |
| Age | | | | | 0.643 |
| <65y | 40(44.4) | 14(51.8) | 16(42.1) | 10(40.0) | |
| ≥65y | 50(55.6) | 13(48.2) | 22(57.9) | 15(60.0) | |
| Smoking | | | | | **0.001** |
| Absent | 74(82.2) | 25(92.6) | 28(73.7) | 21(84.0) | |
| Present | 16(17.8) | 2(7.4) | 10(26.3) | 4(16.0) | |

| | | | | | |
|---|---|---|---|---|---|
| AA intake | | | | | **<0.001** |
| Absent | 63(70.0) | 11(40.7) | 36(94.7) | 16(64.0) | |
| Present | 27(30.0) | 16(59.3) | 2(5.3) | 9(36.0) | |
| Sex | | | | | **0.044** |
| Female | 55(61.1) | 21(77.8) | 18(47.4) | 16(64.0) | |
| Male | 35(38.9) | 6(22.2) | 20(52.6) | 9(36.0) | |
| CKD | | | | | **0.001** |
| 1~2 | 38(42.2) | 6(22.2) | 21(55.2) | 11(44.0) | |
| 3 | 38(42.2) | 9(33.3) | 17(44.7) | 12(48.0) | |
| 4~5 | 14(15.6) | 12(44.4) | 0(0.0) | 2(8.0) | |
| Primary tumour location | | | | | 0.081 |
| Pelvis | 58(64.4) | 22(81.5) | 21(55.3) | 15(60.0) | |
| Ureter | 32(35.6) | 5(18.5) | 17(44.7) | 10(40.0) | |
| Multifocality | | | | | **0.011** |
| Absent | 78(86.7) | 19(70.4) | 36(94.7) | 23(92.0) | |
| Present | 12(13.3) | 8(29.6) | 2(5.3) | 2(8.0) | |
| Tumour size | | | | | **0.023** |
| <3 cm | 38(42.2) | 17(63.0) | 11(28.9) | 10(40.0) | |
| ≥3 cm | 52(57.8) | 10(37.0) | 27(71.1) | 15(60.0) | |
| Architecture | | | | | 0.360 |
| Papillary | 64(71.1) | 21(77.8) | 24(63.2) | 19(76.0) | |
| Sessile | 26(28.9) | 6(22.2) | 14(36.8) | 6(24.0) | |
| T stage | | | | | **0.019** |
| Ta, 1 | 43(47.8) | 19(70.3) | 15(39.5) | 9(36.0) | |
| T2, T3&T4 | 47(52.2) | 8(29.6) | 23(60.5) | 16(64.0) | |
| Grade | | | | | 0.453 |
| Low | 25(27.8) | 7(25.9) | 13(34.2) | 5(20.0) | |
| High | 65(72.2) | 20(74.1) | 25(65.8) | 20(80.0) | |
| N stage | | | | | 0.149 |
| N0 or Nx | 83(92.2) | 27(100.0) | 33(86.8) | 23(92.0) | |
| N1~2 | 7(7.8) | 0(0.0) | 5(13.2) | 2(8.0) | |
| LVI | | | | | 0.087 |
| Absent | 82(8.9) | 27(100.0) | 32(84.2) | 23(92.0) | |
| Present | 8(91.1) | 0(0.0) | 6(15.8) | 2(8.0) | |
| Postoperative | | | | | 0.110 |
| chemotherapy | 81 (90.0) | 27 (100.0) | 33 (86.8) | 21 (84.0) | |
| Absent | 9 (10.0) | 0 (0.0) | 5 (13.2) | 4 (16.0) | |
| Present | | | | | |
| Postoperative radiotherapy | | | | | **0.027** |
| Absent | 85 (94.4) | 27 (100.0) | 33 (86.8) | 25 (100.0) | |
| Present | 5 (5.6) | 0 (0.0) | 5 (13.2) | 0 (0.0) | |

Nx: No lymph node dissection was performed; LVI: lymphovascular invasion.

**Table S3. Univariate and multivariate Cox regression analysis predicting cancer-specific survival and metastasis-free survival.**

| Variables | Cancer specific survival | | | | | | Metastasis-free survival | | | | | |
|---|---|---|---|---|---|---|---|---|---|---|---|---|
| | Univariable | | | Multivariable | | | Univariable | | | Multivariable | | |
| | HR | 95% CI | *P* | HR | 95% CI | *P* | HR | 95% CI | *P* | HR | 95% CI | *P* |
| Age (<65 vs ≥65y) | 1.31 | 0.56~3.07 | 0.532 | | | | 1.64 | 0.72~3.74 | 0.236 | | | |
| Smoking (Absent vs Present) | 0.28 | 0.04~2.08 | 0.212 | | | | 0.22 | 0.03~1.59 | 0.132 | | | |
| AA intake (Absent vs Present) | 0.27 | 0.10~0.76 | **0.013** | | | | 0.26 | 0.09~0.70 | 0.008 | | | |
| Gender (Female vs Male) | 0.82 | 0.29~2.29 | 0.701 | | | | 0.70 | 0.28~1.79 | 0.462 | | | |
| CKD (1~3 vs 4~5) | 0.54 | 0.16~1.82 | 0.318 | | | | 0.51 | 0.15~1.70 | 0.271 | | | |
| Location (Pelvis vs Ureter) | 2.69 | 1.16~6.24 | **0.022** | | | | 2.39 | 1.08~5.27 | **0.031** | | | |
| Multifocality (Absent vs Present) | 0.22 | 0.03~1.63 | 0.138 | | | | 0.21 | 0.03~1.54 | 0.124 | | | |
| Tumour size (<3 cm vs ≥3 cm) | 1.23 | 0.52~2.91 | 0.632 | | | | 0.99 | 0.45~2.18 | 0.972 | | | |
| Architecture (Papillary vs Sessile) | 1.97 | 0.75~5.17 | 0.168 | | | | 1.66 | 0.69~4.04 | 0.261 | | | |
| T stage (Ta, 1&T2 vs T3&T4) | 3.52 | 1.45~8.53 | **0.005** | | | | 3.65 | 1.62~8.21 | **0.002** | | | |
| Grade (Low vs High) | 2.02 | 0.68~6.00 | 0.204 | | | | 2.32 | 0.79~6.76 | 0.125 | | | |
| N (N0 or Nx vs N1~2) | 4.970 | 1.409~17.527 | **0.013** | 5.21 | 1.42~19.19 | **0.013** | 7.04 | 2.53~19.56 | **<0.001** | 6.973 | 2.439~19.933 | **<0.001** |
| LVI (Absent vs Present) | 4.401 | 1.236~15.671 | **0.022** | | | | 5.91 | 2.11~16.53 | **0.001** | | | |
| Signature (AA & DSB vs Age) | 4.054 | 1.577~10.420 | **0.004** | 4.174 | 1.60~10.90 | **0.004** | 3.43 | 1.46~8.06 | **0.005** | 3.442 | 1.437~8.245 | **0.006** |

Nx: No lymph node dissection was performed; LVI: lymphovascular invasion.

**Table S4. The clinicopathologic characteristics for the three UTUC patients from the AA subtype.**

| Patient | Age (yr) | Gender | Germline | Tumour ID | Urothelium ID | Surgery | Grade | pT |
|---|---|---|---|---|---|---|---|---|
| **P1** | 64 | Female | P1RC_1 | P1PT1 | P1BU | 2007.10 | HG | pT3 |
| | | | P1RC_2 | P1PT2 | P1UU | 2015.05 | LG | pT1 |
| | | | | P1BT1 | | 2015.05 | LG | pTa |
| **P2** | 68 | Female | P2RC | P2PT | P2UU | 2016.08 | HG | pT1 |
| **P3** | 53 | Female | P3RC | P3UT | P3UU | 2016.09 | HG | pT2 |

RC= renal cortex; PU= Pelvic urothelium; PT = renal pelvis tumour; UU= ureteral urothelium; UT= ureteral tumour; BU= bladder urothelium; BT= bladder tumour; HG = high grade; and LG = low grade.

**Table S5. Frequently mutated genes among the three patients from the AA subtype.** Nonsynonymous SNVs and indels were annotated by polyphen 2.

| Sample | Probably damaging SNVs-affected genes documented in COSMIC | Probably damaging SNVs-affected genes in the KEGG pathway in cancer |
|---|---|---|
| P1BU | ABL1; BCL11A; CAMTA1; EBF1; FANCD2; HOXA13; KAT6B; NIN; NRG1; PDE4DIP; PTPRB; RUNX1T1; STAT5B; STIL; TRRAP | ABL1; ADCY2; ADCY4; BIRC7; DAPK2; FN1; ITGA2; LAMA1; LAMA2; LAMC3; PRKACG; RUNX1T1; RXRA; STAT5B |
| P1UU | AXIN1; BCORL1; CEP89 DDR2; ELN; ESR1; ETV1; FAT1; FAT4; FLT3; FOXL2; HLF; KDM5C LR; P1B; MECOM; MED12; MLLT6; NR4A3; PAX5; PMS1; PREX2; RBM10; SPECC1; STRN; TCF12; TLX3; TRIP11; WRN | ADCY8; AXIN1; COL4A4; DAPK3; DCC; FLT3; FZD2; FZD7; HGF; LAMA2; MECOM; NOS2; PLCB4 |
| P1PT1 | ARID2; CXCR4; GATA3; HOXD13; KAT6B; NBN; TERT; TFE3; AR; PTEN | CXCR4; FZD9; AR; PTEN |
| P1PT2 | ALDH2; ALK; ARHGAP26; ARID1A; BCL9L; BCR; CACNA1D; CCDC6; CDH1; CEP89; DICER1; DNMT3A; EIF4A2; ELL; EP300; ERBB4; EWSR1; FAT1; FAT4; FBXW7; FGFR1; FLT4; GRIN2A; HIP1; KIT; KMT2A; KMT2C; KMT2D; LIFR; LRP1B; MAP3K1; MECOM; MYH11; NCOR2; NFATC2; NTRK3; PAX7; PDGFB; PMS1; PREX2; PTEN; RNF213; ROS1; SDHA; SMAD2; SMARCA4; SMO; SPECC1; SS18L1; STAT6; TCF3; TET1; TFRC; TSC1; WIF1; ZFHX3; ZNF521; CREBBP; FAT4; PALB2; STK11 | ADCY2; ADCY6 ; ARHGEF11; BCR; CCDC6 ; CDH1; COL4A4; CTNNA2; DCC ; EP300; FGF14; FGFR1; FZD10; GLI3; KIT ; LAMA1; LAMA2; LAMA4; MECOM; MMP2; MMP9; MSH3; PDGFB; PRKACB; PTCH2; PTEN; PTGER2; RASGRP3; SMAD2; SMO; SOS2; WNT10B; CREBBP; MMP9; WNT5B |
| P1BT | ALK; ARHGAP26; ARID1A; BCL9L; BCR; CACNA1D; CCDC6; CD74; CDH1; CEP89; DICER1; DNMT3A; EIF4A2; ELL; EP300; ERBB4; FAT1; FAT4; FBXW7; FGFR1; GRIN2A; HIP1; KIT; KMT2A; KMT2C; KMT2D; LIFR; LRP1B;MAP3K1; MECOM; MYH11; NCOR2; NFATC2; PAX7; PDGFB; PMS1; PREX2; RNF213; ROS1; SDHA; SMAD2; SMO; SPECC1; SS18L1; STAT6; TCF3; TET1; TFRC; TSC1; WIF1; ZFHX3; ZNF521; CREBBP; EWSR1; PALB2 | ADCY2; ADCY6 ; ARHGEF11; BCR; CCDC6; CDH1; COL4A4; CTNNA2; DCC; EDNRA; EP300; FGF14; FGFR1; FZD10; GLI3; KIT; LAMA1; LAMA2; LAMA4; MECOM; MMP2; MMP9; MSH3; PDGFB; PRKACB; PTCH2; PTGER2; RASGRP3; SHH; SMAD2; SMO; SOS2; WNT10B; CREBBP; MMP9 |
| P2UU | ARID2; CLIP1; NF1; KMT2A | COL4A1;ADCY6 |
| P2PT | CDK12; CYLD; KMT2D; TNFRSF14; TP63 | FGF1; LAMA2; ROCK2 |

| Sample | Probably damaging SNVs-affected genes documented in COSMIC | Probably damaging SNVs-affected genes in the KEGG pathway in cancer |
|---|---|---|
| P3PU | AFF4; ALK; ARHGEF12; ATRX; AXIN1; BCR; CASP8; CBFA2T3; CBL; DDR2; EPS15; FGFR4; FLT3; KMT2A; KMT2C; KMT2D; KTN1; LRIG3; LRP1B; MED12; MPL; MSH2; MSN; NACA; NCOA2; NONO; NTRK1; NUP214; PCSK7; PTPRB; PTPRT; QKI; ROS1; SEPT6; SETD2 ; SMAD2; TCF12; TFRC; RRAP; TSHR; WIF1; ZNF521 | ARHGEF12; AXIN1; BCR; CASP8; CBL; DCC; FGF17; FGF5; FLT3 GNGT2; LAMA1; LAMA4; MSH2; NTRK1; PLCB3; RALA; RASGRP4; SMAD2;WNT1; WNT10B; WNT8A |
| P3UT | ABL2; AFF3; APC; AR; ATRX; BCOR; BCORL1; C2orf44; CACNA1D; CCDC6; CHD4; CLIP1; CLTCL1; CREB3L1; CREBBP; CSF3R; CUX1; EBF1; EGFR; ELK4; ERG; FBXW7; FLT3; GMPS; HNF1A; HOOK3; KDM5C; KIAA1549; KMT2C; KMT2D; KTN1; LCP1; LRP1B; MDM2; MSI2; MTOR; NFATC2; NONO; NTRK3; OLIG2; PCM1; PDE4DIP; PDGFB; PER1; PHF6; PICALM; PTPRK; RET; SDHD; SETBP1; SFRP4; SMAD3; SMARCA4 SS18L1; TET1; TLX1; TOP1; TP63; TRRAP; USP6; ATRX; CASC5; CCND2; CREBBP; CUX1; DICER1; EML4; EP300; FOXO4; GAS7; GRIN2A; HIP1; KDM6A; KMT2A; LPP ; MAP3K1; PPM1D; QKI; RUNX1T1; SS18 SUZ12; TPM3 | ADCY2; ADCY4; APC; AR; CCDC6; COL4A6; CREBBP; CSF1R; CSF3R; CTNNA2; EGFR; F2RL3; FASLG; FGF21; FLT3; FZD9; GLI2; HGF; ITGA2B; KIF7; KLK3; LAMA3; LAMA5; LAMC3; MDM2; MMP9; MTOR; PDGFB; PLCB1; PLCB4; PRKCB; PRKCG; PTK2; RASGRP3; RET; ROCK2; SMAD3; STK4; APC2; CREBBP; CTNNA2; DAPK2; EP300; FGF18; GLI3; GNAI1; LAMA5; LPAR1; PIK3CB; RUNX1T1; TPM3 |

**Table S6. Proportion of stromal TIMCs and the number of $CD3^+$ lymphocytes in 76 UTUC patients**

| Variable | Number (%) |
|---|---|
| **Total** | 76 |
| **Stromal of TIMC (%)** | |
| **Focal (0~10)** | 22 (28.9) |
| **Mild (11~30)** | 18 (23.7) |
| **Moderate (31~60)** | 17 (22.4) |
| **Severe (61~100)** | 19 (25.0) |
| **No. of $CD3^+$ TIMC (high power field)** | |
| **0~10** | 24 (31.6) |
| **11~30** | 16 (21.0) |
| **31~60** | 23 (30.3) |
| **61~100** | 13 (17.1) |

**Table S7. The detail list of AA and/or similar compounds-containing herb drugs surveyed by current study.**

| # | Drug names(In Mandarin Chinese) | AA and their derivatives (In Mandarin Chinese) | AA and their derivatives containing herbs | Status |
|---|---|---|---|---|
| 1 | *Long Dan Xie Gan Wan (watered pill,decoction,granule)* | *Guan Mu Tong* | *Aristolochia manshuriensis Kom* | Replaced by *Akebiae Caulis* since 2003 |
| 2 | *Guan Mu Tong Tang Ji (decoction)* | *Guan Mu Tong* | *Aristolochia manshuriensis Kom* | Replaced by *Akebiae Caulis* since 2003 |
| 3 | *Dao Chi Wan (pill)* | *Guan Mu Tong* | *Aristolochia manshuriensis Kom* | Replaced by *Akebiae Caulis* since 2003 |
| 4 | *Gan Lu Xiao Dou Wan (pill)* | *Guan Mu Tong* | *Aristolochia manshuriensis Kom* | Replaced by *Akebiae Caulis* since 2003 |
| 5 | *Dou Long Wan (pill)* | *Guan Mu Tong* | *Aristolochia manshuriensis Kom* | Replaced by *Akebiae Caulis* since 2003 |
| 6 | *Fu Shen Ning Pian (pill)* | *Guang Fang Ji* | *Radix Aristolochiae Fangchi* | Replaced by *Radix Stephaniae Tetrandrae* since 2004 |
| 7 | *Feng Shi Ling Xian Ye (solution)* | *Guang Fang Ji* | *Radix Aristolochiae Fangchi* | Replacedby *Radix Stephaniae Tetrandrae* since 2004 |
| 8 | *Fu Fang Xia Tian Wu Pian (pill)* | *Guang Fang Ji* | *Radix Aristolochiae Fangchi* | Replaced by *Radix Stephaniae Tetrandrae* since 2004 |
| 9 | *Gu Xian Pill (pill)* | *Guang Fang Ji* | *Radix Aristolochiae Fangchi* | Replaced by *Radix Stephaniae Tetrandrae* since 2004 |
| 10 | *Guan Xin Su He Wan (pill,capsule)* | *Qing Mu Xiang* | *Radix Aristolochiae* | Replaced by *Inulae Radix* since 2004 |
| 11 | *Shu Gan Li Qi Wan (pill)* | *Qing Mu Xiang* | *Radix Aristolochiae* | Replaced by *Inulae Radix* since 2004 |
| 12 | *Tong Di Jiao Nang (capsule)* | *Da Qing Mu Xiang* | *Aristolochia austroszechuanica* | Replaced by *Inulae Radixsince* since 2004 |
| 13 | *Xiao Qing Long Tang (decoction)* | *Xi Xin* | *Asari radix et rhizoma* | Available |
| 14 | *Jiu Wei Qiang Huo Tang (decoction)* | *Xi Xin* | *Asari radix et rhizoma* | Available |
| 15 | *Dang Gui Si Ni Tang (decoction)* | *Xi Xin* | *Asari radix et rhizoma* | Available |
| 16 | *Ma Huang Fu Zi Xi Xin Tang (decoction)* | *Xi Xin* | *Asari radix et rhizoma* | Available |
| 17 | *Ke Su Tan Chuan Wan (pill)* | *Ma Dou Ling* | *Fructus Aristolochiae* | Prescription Medicine |
| 18 | *Fu Fang She Dan Chuan Bei San(granule)* | *Ma Dou Ling* | *Fructus Aristolochiae* | Prescription Medicine |
| 19 | *Jing Zhi Ke Su Tan Chuan Wan (pill)* | *Ma Dou Ling* | *Fructus Aristolochiae* | Prescription Medicine |
| 20 | *Wei Fu Ke Li (granule)* | *Ma Dou Ling* | *Fructus Aristolochiae* | Prescription Medicine |
| 21 | *Zhi Ke Hua Tan Wan (pill)* | *Ma Dou Ling* | *Fructus Aristolochiae* | Prescription Medicine |
| 22 | *Chuan Xi Ling Jiao Nang (capsule)* | *Ma Dou Ling* | *Fructus Aristolochiae* | Prescription Medicine |
| 23 | *Fei An Pian (pill)* | *Ma Dou Ling* | *Fructus Aristolochiae* | Prescription Medicine |
| 24 | *Ji Ming Wan (pill)* | *Ma Dou Ling* | *Fructus Aristolochiae* | Prescription Medicine |
| 25 | *Ji Su Wan (pill)* | *Ma Dou Ling* | *Fructus Aristolochiae* | Prescription Medicine |

| # | Drug names(In Mandarin Chinese) | AA and their derivatives (In Mandarin Chinese) | AA and their derivatives containing herbs | Status |
|---|---|---|---|---|
| 26 | *Qi Shi Wei Song Shi Wan (pill)* | *Ma Dou Ling* | *Fructus Aristolochiae* | Prescription Medicine |
| 27 | *Qing Guo Zhi Su Wan (pill)* | *Ma Dou Ling* | *Fructus Aristolochiae* | Prescription Medicine |
| 28 | *Run Fei Hua Tan Wan (pill)* | *Ma Dou Ling* | *Fructus Aristolochiae* | Prescription Medicine |
| 29 | *Shi San Wei Shu Gan Jiao Nang (capsule)* | *Ma Dou Ling* | *Fructus Aristolochiae* | Prescription Medicine |
| 30 | *Zhi Su Hua Tan Wan (pill)* | *Ma Dou Ling* | *Fructus Aristolochiae* | Prescription Medicine |
| 31 | *Xiao Ke Ping Chuan Kou Fu Ye (solution)* | *Ma Dou Ling* | *Fructus Aristolochiae* | Prescription Medicine |
| 32 | *Qing Guo Zhi Ke Wan (pill)* | *Ma Dou Ling* | *Fructus Aristolochiae* | Prescription Medicine |
| 33 | *Xin Bi Tao Xian Pian (pill)* | *Ma Dou Ling* | *Fructus Aristolochiae* | Prescription Medicine |
| 34 | *Zhi Ke Qing Guo Pian (pill)* | *Ma Dou Ling* | *Fructus Aristolochiae* | Prescription Medicine |
| 35 | *Zhi Su Hua Tan Jiao Nang (capsule)* | *Mi Ma Dou Ling* | *Fructus Aristolochiae* | Prescription Medicine |
| 36 | *Dou Shi Jiu Wei Neng Xiao San(granule)* | *Mu Xiang Ma Dou Ling* | *Aristolochia moupinensis* | Prescription Medicine |
| 37 | *Dou Shi Wu Wei Lv Rong Gao Jiao Nang (capsule)* | *Mu Xiang Ma Dou Ling* | *Aristolochia moupinensis* | Prescription Medicine |
| 38 | *Dou Shi Wu Wei Lv Rong Gao Wan (pill)* | *Mu Xiang Ma Dou Ling* | *Aristolochia moupinensis* | Prescription Medicine |
| 39 | *Dou Shi Wu Wei Song Shi Wan (pill)* | *Mu Xiang Ma Dou Ling* | *Aristolochia moupinensis* | Prescription Medicine |
| 40 | *Dou Shi Wu Wei Yu Gan Zi Wan (pill)* | *Mu Xiang Ma Dou Ling* | *Aristolochia moupinensis* | Prescription Medicine |
| 41 | *Dou Shi Wu Wei Zhu Huang San (granule)* | *Mu Xiang Ma Dou Ling* | *Aristolochia moupinensis* | Prescription Medicine |
| 42 | *Feng Shi Sai Long Jiao Nang (capsule)* | *Mu Xiang Ma Dou Ling* | *Aristolochia moupinensis* | Prescription Medicine |
| 43 | *Feng Shi Zhi Tong Wan (pill)* | *Mu Xiang Ma Dou Ling* | *Aristolochia moupinensis* | Prescription Medicine |
| 44 | *Gan Chang Jiao Nang (capsule)* | *Mu Xiang Ma Dou Ling* | *Aristolochia moupinensis* | Prescription Medicine |
| 45 | *Jiu Wei Niu Huang Wan (pill)* | *Mu Xiang Ma Dou Ling* | *Aristolochia moupinensis* | Prescription Medicine |
| 46 | *Qi Wei Hong Hua Shu Sheng San (granule)* | *Mu Xiang Ma Dou Ling* | *Aristolochia moupinensis* | Prescription Medicine |
| 47 | *Qi Wei Hong Hua Shu Sheng Wan (pill)* | *Mu Xiang Ma Dou Ling* | *Aristolochia moupinensis* | Prescription Medicine |
| 48 | *Qing Fei Zhi Ke Wan (pill)* | *Mu Xiang Ma Dou Ling* | *Aristolochia moupinensis* | Prescription Medicine |
| 49 | *Si Wei Zhi Xie Mu Tang San (granule)* | *Mu Xiang Ma Dou Ling* | *Aristolochia moupinensis* | Prescription Medicine |
| 50 | *Wu Wei Zha Xun Wan (pill)* | *Mu Xiang Ma Dou Ling* | *Aristolochia moupinensis* | Prescription Medicine |
| 51 | *Tian Xian Teng San (granule)* | *Tian Xian Teng* | *Herba Aristolochiae* | Prescription Medicine |
| 52 | *He Wei Jiang Ni Jiao Nang (capsule)* | *Tian Xian Teng* | *Herba Aristolochiae* | Prescription Medicine |
| 53 | *Xiang Teng Jiao Nang (capsule)* | *Tian Xian Teng* | *Herba Aristolochiae* | Prescription Medicine |
| 54 | *Yun Xue Tang (decoction)* | *Tian Xian Teng* | *Herba Aristolochiae* | Prescription Medicine |
| 55 | *Fu Fang Feng Shi Yao Jiu (medicinal liquor)* | *Xun Gu Feng* | *herba Aristolochiae mollissimae* | Prescription Medicine |

| # | Drug names(In Mandarin Chinese) | AA and their derivatives (In Mandarin Chinese) | AA and their derivatives containing herbs | Status |
|---|---|---|---|---|
| 56 | *Fu Fang Quan Can Pian (pill)* | *Xun Gu Feng* | *herba Aristolochiae mollissimae* | Prescription Medicine |
| 57 | *Qu Feng Chu Shi Yao Jiu (medicinal liquor)* | *Xun Gu Feng* | *herba Aristolochiae mollissimae* | Prescription Medicine |
| 58 | *San She Yao Jiu (medicinal liquor)* | *Xun Gu Feng* | *herba Aristolochiae mollissimae* | Prescription Medicine |
| 59 | *Shen Nong Yao Jiu (medicinal liquor)* | *Xun Gu Feng* | *herba Aristolochiae mollissimae* | Prescription Medicine |
| 60 | *Yi Shen Juan Bi Wan (pill)* | *Xun Gu Feng* | *herba Aristolochiae mollissimae* | Prescription Medicine |
| 61 | *Dou Zhong Zhuang Gu Jiao Nang (decoction)* | *Xun Gu Feng* | *herba Aristolochiae mollissimae* | Prescription Medicine |
| 62 | *Dou Zhong Zhuang Gu Wan (pill)* | *Xun Gu Feng* | *herba Aristolochiae mollissimae* | Prescription Medicine |
| 63 | *Feng Shi Ning Yao Jiu (medicinal liquor)* | *Xun Gu Feng* | *herba Aristolochiae mollissimae* | Prescription Medicine |
| 64 | *Shao Lin Zheng Gu Jing (pill)* | *Xun Gu Feng* | *herba Aristolochiae mollissimae* | Prescription Medicine |
| 65 | *Yi Shen Juan Bi Wan (pill)* | *Xun Gu Feng* | *herba Aristolochiae mollissimae* | Prescription Medicine |
| 66 | *Fu Fang Wei Tong Jiao Nang (capsule)* | *Zhu Sha Lian* | *Aristolochia cinnabarina* | Prescription Medicine |
| 67 | *Jiu Long Jie Dou Jiao Nang (capsule)* | *Zhu Sha Lian* | *Aristolochia cinnabarina* | Prescription Medicine |
| 68 | *Bao Wei Jiao Nang (capsule)* | *Zhu Sha Lian* | *Aristolochia cinnabarina* | Prescription Medicine |
| 69 | *Jin Zhu Zhi Xie Pian (pill)* | *Zhu Sha Lian* | *Aristolochia cinnabarina* | Prescription Medicine |
| 70 | *Zhu Sha Lian Jiao Nang (capsule)* | *Zhu Sha Lian* | *Aristolochia cinnabarina* | Prescription Medicine |
